# RNA Small-Molecule Binding-Site Prediction by Integrating RNA Language Models and Structural Geometry

**DOI:** 10.64898/2026.11.10.765632

**Authors:** Nandita Sharma, Pralay Mitra

## Abstract

RNAs are emerging as promising therapeutic targets, and small-molecule binding can modulate their structure and function, creating opportunities for therapeutic development. Accurate identification of RNA–small-molecule binding sites is therefore important for understanding RNA–ligand interactions and facilitating RNA-targeted drug discovery. However, the structural complexity of RNA and the limited availability of experimentally characterized RNA–ligand complexes pose challenges for computational binding-site prediction. In this study, we propose RSMBind, a graph attention-based deep learning framework for nucleotide-level prediction of RNA–small-molecule binding sites. The model integrates geometric descriptors of the local three-dimensional environment with pretrained RNA language model embeddings to capture complementary structural and sequence information. We investigate three pretrained RNA language models—RNA-BERT, ERNIE-RNA and RNA-FM—and assess the contributions of sequence representations and geometric features through feature-level ablation experiments. The model is developed using the TR60 and HARIBOSS datasets and evaluated on four independent benchmark datasets: RB9, TE18, TL12 and JL10. RSMBind achieves the highest MCC among the compared methods on three of the four benchmarks, with relative improvements of 17.1% on RB9, 7.2% on TL12 and 16.0% on JL10 over the best previously reported results, while achieving competitive performance on TE18. Furthermore, RSMBind retains predictive capability when using only pretrained RNA language model embeddings, highlighting their potential utility when explicit tertiary structural features are unavailable. Structural analysis indicates differences in prediction performance across RNA structural regions, with loop-associated binding sites generally being captured more effectively than those embedded within stem regions. Additionally, 100 ns molecular dynamics simulations of selected RNA–ligand complexes provide insights into the relationship between predicted binding probabilities and dynamic ligand proximity, further highlighting the role of conformational dynamics in evaluating structure-based predictions. The source code is available at [ https://github.com/sharma-nandita/RSMBind].

## 1. Introduction

RNA has emerged as an important therapeutic target because of its diverse regulatory and functional roles. Many non-coding RNAs participate in cellular regulation, and their dysregulation has been associated with several diseases, including cancer and neurodegenerative disorders [1] [2] . Small molecules that bind RNA can modulate its function, highlighting the potential of RNA-targeted therapeutics. These RNA molecules adopt complex secondary and tertiary structures that create distinct binding environments for small molecules [3] . Identifying the nucleotides involved in ligand binding is therefore an important step toward understanding RNA–ligand interactions and facilitating the development of RNA-targeted therapeutics [4]. Accurate identification of RNA–small-molecule binding sites remains challenging, while experimental approaches are often time-consuming and resource-intensive. This has motivated the development of computational methods for efficient binding-site prediction [5]. Early methods, such as Rsite [6] and Rsite2 [7] used distance-based relationships derived from tertiary and secondary RNA structures, respectively. RBind[8] represented RNA as a nucleotide interaction network and used graph-theoretical properties to identify potential binding sites. However, its use of predefined thresholds can favor high precision at the expense of sensitivity, limiting its ability to identify a broader range of binding nucleotides. RNAsite[9] combined evolutionary conservation and structural features, including solvent accessibility and graph-based properties, using a Random Forest classifier. Although it demonstrated the value of integrating sequence and structural information, its reliance on handcrafted features may limit its ability to capture complex nucleotide interactions. Deep learning methods have enabled more effective learning of complex relationships between RNA sequence and structural features. RLBind [10] combined global RNA information with local nucleotide features using a convolutional neural network, capturing both local and long-range interactions. It can also use modeled structures when experimental tertiary structures are unavailable. ZeSTa[11] integrated sequence, structural, geometric, surface, and network features using ResNet18 to predict binding sites. RNet[12] used distance-based dynamic graphs to represent RNA tertiary interactions and incorporated graph features into an ensemble learning framework for binding-site prediction. Further Graph-based methods have been combined with pretrained RNA language models to integrate sequence and structural information for binding-site prediction. RNABind [13] combines pretrained RNA language model embeddings with an Equivariant Graph Convolutional Layer (EGCL) to capture both contextual sequence information and spatial relationships between nucleotides. RLsite [14] employs a Graph Attention Network (GAT) to incorporate RNA secondary- and tertiary-structure information for binding-site prediction, while also utilizing pretrained ERNIE-RNA-derived sequence embeddings as input to a multilayer perceptron (MLP) for prediction. MVRBind [15] employs multi-layer graph convolutions with multi-scale edges and transformer-based representations to capture nucleotide dependencies. GATRsite [16] also combines pretrained RNA language model representations with graph attention networks to jointly represent sequence and structural information. Further In CoBRA [17] binding-sites are predicted using RiNALMo-derived sequence representations only, with a relatively simple MLP-based classifier. More recently, GRASSP [18] combined ERNIE-RNA-derived sequence embeddings and predicted secondary-structure features using a two-step graph-attention refinement network with adaptive gating. These methods show different strategies for combining pretrained RNA representations with structural information for binding-site prediction.

Despite recent advances, RNA–small-molecule binding-site prediction remains challenging because binding nucleotides can occur in both paired and unpaired regions and in spatially complex structural environments. Limited availability of experimentally characterized RNA– ligand complexes further constrains model development. To address these challenges, we propose a graph attention-based framework for nucleotide-level RNA–small-molecule binding-site prediction. RNA molecules are represented as graphs, with nucleotides as nodes and three types of edges representing backbone connectivity, secondary-structure base-pairing relationships, and spatial proximity. Spatial edges connect nucleotides within an 8 Å cutoff and incorporate inter-nucleotide distance information. The framework combines geometry-based descriptors of the local three-dimensional environment with pretrained RNA language model embeddings. We investigate three pretrained models, RNA-BERT, ERNIE-RNA, and RNA-FM, to assess their contribution to binding-site prediction. A single-layer graph attention network with a residual connection integrates information from neighboring nucleotides while maintaining a relatively simple architecture suited to the limited availability of training data. We evaluate the proposed framework on four independent benchmark datasets: RB9, TE18, TL12, and JL10. The results demonstrate superior overall performance on three datasets and competitive performance on TE18. Feature-level ablation experiments further examine the contributions of pretrained sequence representations and geometry-based descriptors. Notably, the framework retains promising predictive performance when using only language model-derived sequence embeddings, indicating its potential applicability when experimentally determined tertiary structures are unavailable. In addition, we analyze prediction performance across RNA structural regions and conduct molecular dynamics (MD) simulations of selected RNA–ligand complexes for 100 ns. These case studies examine the relationship between model predictions and ligand proximity over the simulation trajectory, providing insights into prediction errors and the structural behavior of RNA–ligand interactions.

## 2. Materials and Methods

### 2.1. Dataset Preparation

RNA–small-molecule binding site prediction is formulated as a nucleotide-level binary classification problem, in which each nucleotide is classified as binding or non-binding. In this study, small molecules are defined as compounds with molecular weights below 1000 Da, including metal ions, natural metabolites, and drug-like compounds. A nucleotide was labelled as binding if the minimum distance between any heavy atom of the nucleotide and any heavy atom of the associated ligand was ≤4 Å. This criterion is consistent with previous RNA–ligand binding site prediction studies. Here six publicly available datasets are considered: TR60, HARIBOSS, TE18, RB9, TL12, and JL10. TR60 and HARIBOSS are combined to construct the development dataset, whereas TE18, RB9, TL12, and JL10 are reserved for independent evaluation. All RNA–ligand structures are obtained from the Protein Data Bank (PDB)[19].

The TR60 and TE18 datasets originated from the RNAsite study [9], which collected 712 RNA–small-molecule complexes from the PDB. Water molecules and crystallization additives were excluded, and RNA chains containing 20–1500 nucleotides were considered. For complexes containing multiple ligands, the binding sites associated with each ligand were retained. Structural redundancy was reduced using TM-scoreRNA with a cutoff of 0.3. When two RNA chains exceeded this threshold, the chain with more binding nucleotides was retained. The resulting 78 representative chains were further clustered at a 30% sequence-similarity threshold, producing 57 clusters. Of these chains, 60 were assigned to TR60 and 18 to TE18. TE18 was retained as an independent test set and was not used during model development. The HARIBOSS dataset was obtained from the work of [20], who collected 862 RNA–small-molecule structures from the PDB and filtered them based on sequence and structural similarity. In our study, HARIBOSS dataset is combined with TR60 dataset to increase the diversity of the development dataset, following previous RNA binding site prediction studies [17] [18]. For the combined dataset, RNA–ligand contacts were identified using the same 4 Å heavy-atom distance criterion. Water, sulfate, phosphate, glycerol, ethylene glycol, and polyethylene glycol are excluded from ligand assignment because these molecules commonly occur as crystallization additives or cryoprotectants. RNA chains with incomplete coordinates required for three-dimensional feature extraction and those longer than 438 nucleotides are excluded. After ligand assignment and quality filtering, 358 RNA chains containing 24,107 nucleotides are retained for model development. Their lengths ranged from 20 to 418 nucleotides, with a mean of 67.34 ± 55.43 and a median of 53 nucleotides. Of the 24,107 nucleotides, 6,870 (28.50%) were labelled as binding and 17,237 (71.50%) as non-binding, giving a positive-to-negative ratio of approximately 1:2.51. The development dataset was evaluated using stratified five-fold cross-validation. In each fold, the data were divided into training and validation subsets while preserving the class distribution as closely as possible. Five models were trained using the respective training folds. Each trained model was evaluated on the independent test datasets, and the predictions from the five models were averaged to obtain the final ensemble prediction for each nucleotide.

Four datasets, RB9, TE18, TL12, and JL10, were used only for external evaluation and were excluded from model training and parameter selection. RB9 was derived from a previously published benchmark used by RBind[8] and RNAsite[9] . The original RB19 dataset contained structures overlapping with TR60. Ten overlapping structures were removed to reduce data leakage, leaving nine non-redundant RNA chains in RB9. TE18 contains 18 RNA chains selected during the construction of the original RNAsite benchmark, as described above. It was retained for independent evaluation and was not used for model development. TL12 was derived from the benchmark introduced in [11] and contains RNA structures with loops but without junction loops. It provides an evaluation set representing RNA architectures without junction-loop complexity. JL10 was constructed to evaluate model performance on structurally complex RNAs containing junction loops. RNA structures deposited in the PDB after January 2021 were considered to reduce overlap with earlier benchmarks [11]. Additional filtering was applied to remove structures with excessive numbers of junction loops, low proportions of binding nucleotides, or high sequence redundancy. The resulting dataset contains 10 RNA chains. Together, the four independent test sets provide complementary evaluations across RNA structures with different sequence characteristics, binding-site distributions, and structural complexity. The dataset statistics are summarized in Table 1, and detailed information for each test set is provided in Supplementary Tables S1– S4.

**Table 1.** Summary of the four independent benchmark datasets used for evaluating the model.

| **Dataset** | **RNAs** | **Total nts** | **Positive** | **Positive (%)** | **Negative** | **Pos: Neg ratio** | **Length (mean ± SD)** |
|---|---|---|---|---|---|---|---|
| RB9 | 9 | 459 | 133 | 28.98 % | 326 | 1:2.45 | 51.00 ± 29.03 |
| TE18 | 18 | 558 | 187 | 33.51 % | 371 | 1:1.98 | 34.88 ± 17.49 |
| TL12 | 12 | 516 | 179 | 34.69 % | 337 | 1:1.88 | 43.00 ± 17.49 |
| JL10 | 10 | 827 | 267 | 32.29 % | 560 | 1:2.10 | 82.70 ± 20.47 |

### 2.2. Feature Construction

To represent the sequence, surface, and three-dimensional structural properties of RNA, here we constructed a multimodal nucleotide-level feature representation. Each nucleotide was represented using a combination of handcrafted structural features and pretrained RNA language model embeddings. The resulting feature vector was assigned to the corresponding nucleotide node in the RNA graph. The handcrafted features characterize the local structural environment, whereas the language model embeddings capture contextual sequence information. These complementary representations were integrated before graph-based learning.

#### 2.2.1 Handcrafted Structural Features

Nine handcrafted features were extracted to characterize the spatial environment, solvent accessibility, and graph topology of each nucleotide. Five spatial descriptors were calculated from the three-dimensional RNA coordinates: distance from the RNA centroid, normalized sequence position, number of neighboring nucleotides within an 8 Å cutoff, and the mean and minimum distances to spatial neighbors. The centroid distance describes the nucleotide’s position relative to the RNA’s geometric center, while normalized sequence position represents its relative location along the sequence. The neighborhood-based descriptors characterize local structural density and packing. For the mean and minimum neighbor-distance calculations, immediate sequence neighbors were excluded when non-backbone spatial contacts were available, reducing the influence of covalent backbone contacts. The neighbor-count feature included all nucleotides within the 8 Å cutoff.

Two solvent-accessibility descriptors were calculated using FreeSASA [21] tool. The first was the SASA of each nucleotide normalized by the maximum nucleotide SASA within the corresponding RNA structure. The second was the standardized SASA, calculated by subtracting the mean and dividing by the standard deviation of nucleotide SASA values within that structure. These descriptors characterize the relative exposure and burial of nucleotides. Further, two graph-topological descriptors were also calculated from the RNA graph: node degree and local clustering coefficient. Node degree represents the number of unique neighboring nucleotides connected to a given nucleotide through the constructed RNA graph. It therefore reflects the local connectivity of a nucleotide considering backbone, base-pair, and spatial interactions. The clustering coefficient measures the extent to which these neighbors are interconnected. Together, these descriptors formed a nine-dimensional handcrafted feature vector for each nucleotide. The distributions of binding and non-binding nucleotides were examined to characterize the discriminative properties of these features (Supplementary Figure S1). Their individual contributions were further assessed through feature ablation experiments, as described in the Results section.

#### 2.2.2 Pretrained RNA language model embeddings

To capture contextual sequence information that may not be fully represented by the handcrafted structural descriptors, pretrained RNA language model embeddings were incorporated into the nucleotide-level representations. Three pretrained RNA language models were considered: RNABERT, ERNIE-RNA, and RNA-FM. These models were selected because they were pretrained on large collections of non-coding RNA sequences and have been used in previous RNA structure and interaction prediction studies. RNABERT[21] was introduced as a pretrained Transformer-based model for learning contextual representations of RNA sequences. It produces a 120-dimensional representation for each nucleotide. ERNIE-RNA[22] employs a 12-layer Transformer architecture and was pretrained using masked language modeling on large-scale non-coding RNA sequences, incorporating sequence and structural information during pretraining. It produces a 768-dimensional representation for each nucleotide. RNA-FM[23] is a bidirectional Transformer-based RNA foundation model comprising 12 Transformer encoder layers and was pretrained on approximately 23.7 million non-coding RNA sequences. It generates a 640-dimensional representation for each nucleotide. Thus, the three language models produce nucleotide-level embeddings of substantially different dimensionalities:

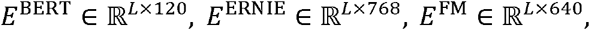

Where *L* denotes the RNA sequence length. Because the original embedding dimensions are substantially larger than the handcrafted feature representation, directly concatenating the embeddings would give the high-dimensional language-model features disproportionate influence in the subsequent feature-fusion stage. Therefore, each embedding type was independently projected into a lower-dimensional latent space before fusion. For RNABERT, the 120-dimensional representation was projected to 32 dimensions. ERNIE-RNA and RNA-FM representations were independently projected from 768 and 640 dimensions to 128 dimensions, respectively. The projection networks consisted of layer normalization and learnable fully connected nonlinear transformations. For an input embedding *e*_*i*_, the corresponding projection can be represented generally as

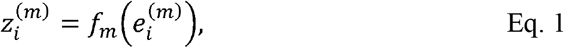

Where *m*denotes the selected language model and *f*_*m*_ (·) is its learnable projection network. The projection networks were trained jointly with the downstream graph neural network, while the parameters of the pretrained language models themselves remained frozen. The resulting projected language-model representations were concatenated with the handcrafted nucleotide-level features. For a nucleotide *i*, the fused representation can therefore be expressed as

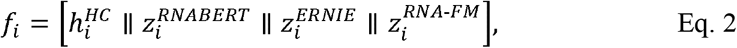

where 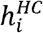 denotes the handcrafted feature vector and only the language-model representations included in a particular experimental configuration are concatenated. The dimensionality of the fused representation therefore depends on the selected feature configuration. This fused representation is subsequently transformed into the common 128-dimensional hidden space by the input projection layer before graph message passing. These configurations were evaluated to investigate the individual and combined contributions of the pretrained representations to RNA–small-molecule binding-site prediction.

### 2.3 RNA graph construction

RNA tertiary structures were represented as graphs to model interactions between nucleotides. For an RNA molecule containing *N* nucleotides, we constructed a graph

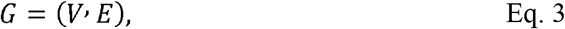

where each node *v*_*i*_ ∈ *V* represents one nucleotide and each edge *e*_*ij*_ ∈ *E* represents a structural or spatial relationship between two nucleotides. Each nucleotide node was associated with a feature vector containing the corresponding nucleotide-level handcrafted features and, when enabled, the projected representations derived from pretrained RNA language models. These feature components were concatenated and transformed into a common 128-dimensional representation before being provided to the GATv2 layer. Thus, the graph structure and node features jointly encode the local biochemical characteristics and contextual sequence information of each nucleotide.

Three types of edges were considered: backbone edges, base-pair edges, and spatial proximity edges.

- *Backbone edges* connect consecutive nucleotides along the RNA sequence. These edges represent the covalent connectivity of the RNA backbone. Bidirectional edges were used so that information could propagate in both directions along the sequence.
- *Base-pair edges* represent nucleotide pairing relationships obtained from the secondary-structure dot-bracket representation. A bidirectional edge was introduced between two nucleotides forming a base pair. These edges provide structural information that is not captured by sequence adjacency alone.
- *Spatial edges* were introduced between two nucleotides whose three-dimensional distance was less than 8 Å. The Euclidean distance between nucleotide coordinates was used to determine spatial proximity. For a pair of nucleotides *i* and *j*, the spatial distance was calculated as

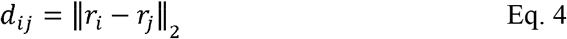

where *r*_*i*_ and *r*_*j*_ denote the three-dimensional coordinates associated with nucleotides *i* and *j*, respectively. A spatial edge was created when *d*_*ij*_ < 8 Å. Then the spatial distance was normalized by the 8 Å cutoff:

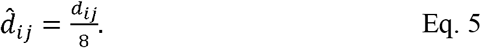

Each edge was assigned an edge-type indicator distinguishing backbone, base-pair, and spatial interactions. The resulting edge attribute was represented as a four-dimensional vector,

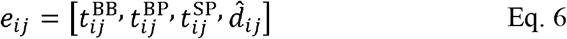

where 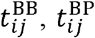 and 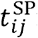 indicate the backbone, base-pair, and spatial edge types, respectively, and 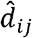 denotes the normalized spatial distance. For backbone and base-pair edges, the distance component was set to zero, whereas spatial edges additionally carried the normalized Euclidean distance. The resulting graph integrates sequence connectivity, secondary-structure interactions, and three-dimensional spatial proximity within a single graph representation. These node and edge attributes are subsequently provided to the edge-aware GATv2 layer, enabling message passing to account for both nucleotide representations and the structural relationship between connected nucleotides.

### 2.4 GATv2-based Model Architecture

The proposed model predicts RNA–small-molecule binding sites at the nucleotide level by integrating pretrained RNA language-model embeddings, handcrafted nucleotide descriptors, and graph-based structural information. The RNA graph and its edge attributes are constructed as described in Section 2.3, with the four-dimensional edge attribute vector (*e*_*i j*_ ) defined in Eq. (6). The embeddings from RNABERT, RNA-ERNIE, and RNA-FM are processed by separate feed-forward projection networks to obtain 32-, 128-, and 128-dimensional representations, respectively. These representations are concatenated with the nine handcrafted descriptors, and a linear layer projects the combined features into a shared 128-dimensional space. The resulting representation of nucleotide (*i*) is denoted by (*h*_*i*_ ∈ ℝ^128^), and (*H*) denotes the matrix of node representations in the graph (Figure 1a).

**Figure 1.**
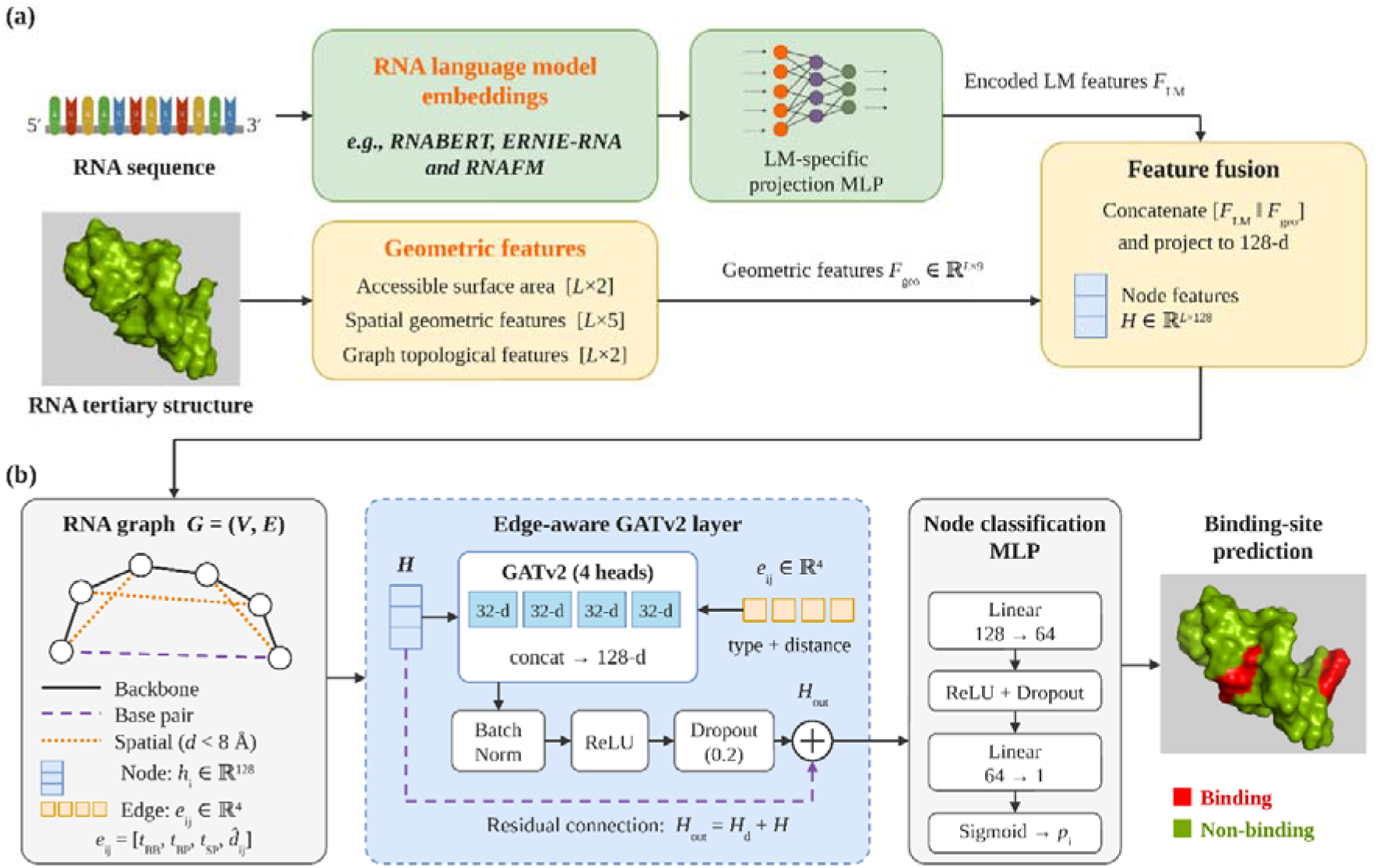
Overview of the proposed RNA ligand-binding site prediction framework. (a) Language-model embeddings from the RNA sequence and geometric features from the tertiary structure are fused into 128-dimensional node features. (b) The RNA graph, with backbone, base-pair and spatial edges, is processed by a single edge-aware GATv2 layer with a residual connection, followed by an MLP classifier that outputs a binding probability for each nucleotide.

The fused node representations are refined using a single GATv2 layer [24], which aggregates information from neighboring nucleotides through attention. GATv2 was selected over the original GAT [25] because its attention mechanism allows the relative importance of a neighbor to depend on the target nucleotide. This is relevant to binding-site prediction, where the contribution of a neighboring nucleotide may vary with the structural context of the nucleotide being classified. For attention head (m), the attention logit between target node (i) and neighbor (j) is computed as follows:

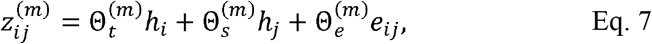

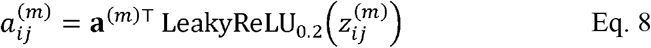

where 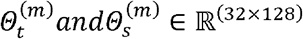 are learnable transformations of the target and neighbor nodes, 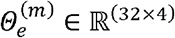 transforms the edge attributes, *a*^(m)^ ∈ ℝ^32^ is a learnable attention vector, and the negative slope of the LeakyReLU is 0.2. The logits are normalized across the neighborhood *N*(*i*) of the target node using a softmax function,

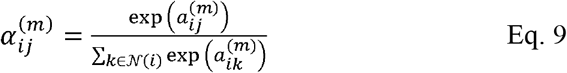

where *N*(*i*) comprises all nodes with an edge directed to *i*, together with *i* itself through a self-loop. The output of head *m* is the attention-weighted sum of the transformed neighbor representations,

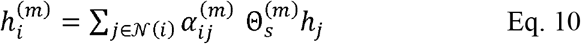

Edge attributes therefore influence the attention coefficients but are not included in the aggregated message. The layer uses four attention heads of 32 dimensions each, and their outputs are concatenated to restore the 128-dimensional representation,

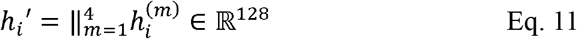

During training, dropout with a rate of 0.1 is applied to the attention coefficients 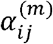.

Because the input and output of the GATv2 layer have the same dimensionality, a residual connection can be applied without an additional projection. The graph-attention output is batch-normalized, passed through a ReLU activation, and regularized with dropout (rate 0.2), and the result is added to the layer input,

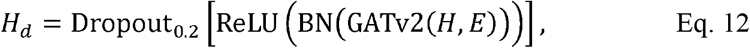

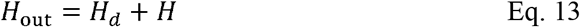

where E denotes the edge indices and edge attributes of the graph. The skip connection preserves the fused nucleotide information, while the GATv2 layer adds neighborhood-dependent structural context (Figure 1b). A single layer restricts message passing to the one-hop neighborhood, which limits over-smoothing from multi-hop propagation, as supported by the ablation experiments in result section.

The refined representation 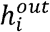 of each nucleotide is finally passed to a two-layer fully connected classifier,

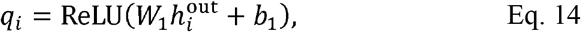

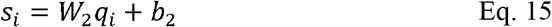

with *W*_1_ ∈ ℝ^(64 ×128)^, *b*_1_ ∈ *ℝ*^64^, *W*_2_ ∈ *ℝ*^(1×64)^, and *b*_2_ ∈ *ℝ*. The logit *s*_*i*_ is converted to a binding probability by the sigmoid function, *p*_*i*_ *= σ*(*s*_*i*_), and nucleotide *i* is assigned to the binding class if *p*_*i*_ exceeds the decision threshold.

### 2.5 Training and Optimization

The model was trained as a nucleotide-level binary classifier using binary cross-entropy with logits as the training objective. To account for the imbalance between ligand-binding and non-binding nucleotides, a positive-class weight of 1.5 was incorporated into the loss function. The model was optimized using the Adam optimizer with a learning rate of 1x 10^−3^and a weight decay of 1x 10^−5^. Training was performed for a maximum of 20 epochs with a batch size of 8. The random seed was fixed to 42 to improve reproducibility. The HARIBOSS + TR60 training set was divided into five folds, and the checkpoint with the highest validation MCC was retained for each fold. For each independent test set, the final nucleotide-level prediction probabilities were obtained by taking the arithmetic mean of the probabilities predicted by the five fold-trained models.

### 2.6 Evaluation Metrics

The performance of RSMBind was evaluated using precision, recall, F1-score, Matthews correlation coefficient (MCC), and the area under the receiver operating characteristic curve (AUROC). Precision measures the proportion of predicted binding nucleotides that are correctly identified, whereas recall measures the proportion of actual binding nucleotides that are correctly identified. They are defined as

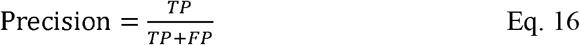

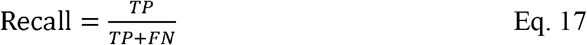

The F1-score is the harmonic mean of precision and recall:

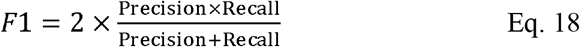

The Matthews correlation coefficient (MCC) evaluates the agreement between predicted and actual classes by incorporating all four confusion-matrix components:

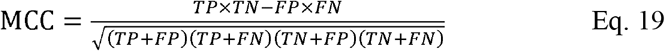

Here, (TP), (TN), (FP), and (FN) denote true positives, true negatives, false positives, and false negatives, respectively. MCC ranges from −1 to 1, where 1 indicates perfect prediction, 0 indicates no correlation between predictions and actual labels, and −1 indicates complete disagreement. AUROC measures the model’s ability to distinguish between binding and non-binding nucleotides across classification thresholds. It is calculated as the area under the receiver operating characteristic (ROC) curve, which plots the true-positive rate against the false-positive rate at varying thresholds. For independent testing, precision, recall, F1-score, and MCC were calculated from the ensemble predictions using a fixed decision threshold of 0.5. AUROC was calculated from the continuous ensemble probabilities without applying a classification threshold.

## 3. Results and Discussion

### 3.1. Comparison with Existing RNA Small-Molecule Binding-Site Prediction Methods

RSMBind was trained on the combined HARIBOSS and TR60 datasets and evaluated on four independent benchmarks (RB9, TE18, TL12, and JL10) after removing overlapping sequences between the training and test sets, following the CoBRA evaluation protocol. No additional training was performed on the test datasets. Performance was compared with previously reported results for Rsite, Rsite2, RNAsite, RBind, RLBind, ZeSTa, RNABind, MVRBind, CoBRA, and GRASSP (Table 2).

**Table 2.** Comparison of RSMBind with existing RNA small-molecule binding-site prediction methods on four independent benchmark datasets. Results for existing methods were obtained from previously published studies, primarily CoBRA and ZeSTa. The highest value for each metric is highlighted in bold. A dash (-) indicates that the corresponding value was not reported.

|  |  | Rsite | Rsite2 | RNAsite | RBind | RLBind | ZeSTa | RNABind | MVRBind | CoBRA | GRASSP | Our Method<br>(**RSMBind**) |
|---|---|---|---|---|---|---|---|---|---|---|---|---|
| **RB9 Test set** | MCC | 0.159 | 0.072 | 0.426 | 0.278 | - | 0.398 | - | - | 0.557 | - | **0.652** |
|  | AUROC | 0.575 | 0.528 | 0.790 | 0.592 | - | 0.786 | 0.865 |  | 0.844 | - | **0.909** |
|  | Precision | 0.430 | 0.382 | 0.750 | 0.681 | - | **0.767** | - |  | 0.750 | - | 0.765 |
|  | Recall | 0.353 | 0.187 | 0.378 | 0.230 | - | 0.408 | - |  | 0.650 | - | **0.7368** |
| **TE18 Test set** | MCC | 0.071 | 0.010 | 0.253 | 0.187 | 0.324 | 0.327 | - | 0.351 | 0.190 | **0.359** | 0.350 |
|  | AUROC | 0.590 | 0.474 | **0.776** | 0.559 | 0.720 | 0.709 | 0.737 | 0.745 | 0.647 | 0.746 | 0.764 |
|  | Precision | 0.449 | 0.370 | 0.675 | 0.655 | 0.681 | **0.729** | - | 0.645 | 0.562 | 0.662 | 0.580 |
|  | Recall | 0.288 | 0.214 | 0.263 | 0.173 | 0.345 | 0.379 | - | 0.342 | 0.357 | 0.425 | **0.544** |
| **TL12 Test set** | MCC | - | - | - | - | - | 0.440 | - |  | 0.546 | 0.611 | **0.655** |
|  | AUROC | - | - | - | - | - | 0.704 | 0.754 |  | 0.816 | **0.874** | 0.869 |
|  | Precision | - | - | - | - | - | 0.740 | - |  | 0.816 | 0.827 | **0.876** |
|  | Recall | - | - | - | - | - | 0.514 | - |  | 0.599 | 0.646 | **0.675** |
| **JL10 Test set** | MCC | 0.046 | 0.007 | - | - | - | 0.211 | - |  | 0.293 | 0.343 | **0.398** |
|  | AUROC | 0.477 | 0.504 | - | - | - | 0.592 | 0.592 |  | 0.739 | 0.760 | **0.768** |
|  | Precision | 0.295 | 0.338 | - | - | - | 0.549 | - |  | 0.582 | 0.587 | **0.661** |
|  | Recall | 0.194 | 0.131 | - | - | - | 0.296 | - |  | **0.531** | 0.486 | 0.475 |

RSMBind achieved MCC values of 0.652, 0.350, 0.655, and 0.398 on RB9, TE18, TL12, and JL10, respectively, with corresponding AUROC values of 0.909, 0.764, 0.869, and 0.768 (Table 2). It achieved the highest reported MCC among the compared methods on RB9, TL12, and JL10. Compared with CoBRA, MCC increased by 17.1%, 20.0%, and 35.8% on these datasets, respectively. RSMBind also outperformed GRASSP in MCC on TL12 and JL10 by 7.2% and 16.0%, respectively. On TE18, RSMBind achieved an MCC of 0.350, comparable to the reported values for MVRBind (0.351) and GRASSP (0.359), and higher than that of CoBRA (0.190). Its AUROC of 0.764 was also higher than those reported for MVRBind (0.745), GRASSP (0.746), and CoBRA (0.647). Overall, RSMBind achieved strong performance on three independent benchmarks and competitive performance on TE18, supporting its effectiveness for ligand-unaware RNA small-molecule binding-site prediction.

### 3.2. Effect of Training-Set Expansion on Generalization

To assess the effect of training-set expansion, we compared RSMBind trained on TR60 alone with the same model trained on the combined TR60+HARIBOSS dataset. Both configurations used the same model architecture and training settings. Performance was evaluated on the four independent benchmark datasets, with results summarised in Table 3.

**Table 3.** Performance of RSMBind on four independent benchmark datasets when trained on TR60 alone and on the combined TR60+HARIBOSS dataset.

| **Training dataset** | **Test set** | **MCC** | **Precision** | **Recall** | **F1 score** |
|---|---|---|---|---|---|
| TR60 | RB9 | 0.520 | 0.705 | 0.594 | 0.645 |
|  | TE18 | 0.305 | 0.605 | 0.404 | 0.485 |
|  | JL10 | 0.254 | 0.603 | 0.285 | 0.387 |
|  | TL12 | 0.248 | 0.609 | 0.357 | 0.449 |
| TR60+HARIBOSS | RB9 | 0.6523 | 0.7656 | 0.7368 | 0.7509 |
|  | TE18 | 0.3500 | 0.5808 | 0.5449 | 0.5623 |
|  | JL10 | 0.3982 | 0.6616 | 0.4756 | 0.5533 |
|  | TL12 | 0.6555 | 0.8760 | 0.6751 | 0.7625 |

Training on TR60+HARIBOSS improved MCC across all four benchmarks, from 0.520 to 0.652 on RB9, 0.305 to 0.350 on TE18, 0.254 to 0.398 on JL10, and 0.248 to 0.655 on TL12. F1-score also increased on every dataset, from 0.645 to 0.751 on RB9, 0.485 to 0.562 on TE18, 0.387 to 0.553 on JL10, and 0.449 to 0.763 on TL12. The largest improvement occurred on TL12, followed by JL10, whereas the gain on TE18 was comparatively modest. These results indicate that incorporating HARIBOSS improved predictive performance across the evaluated benchmarks, although the magnitude of improvement varied by dataset.

### 3.3. Sequence-Derived Representations in the Absence of Tertiary-Structure Features

We investigated whether pretrained RNA language-model (LM) representations could support nucleotide-level binding-site prediction without explicitly provided tertiary-structure information. RNABERT, ERNIE-RNA, and RNA-FM were evaluated individually and in combination using their sequence-derived embeddings of 120, 768, and 640 dimensions, respectively. The embeddings were projected to lower-dimensional representations before integration into the graph-based model. Handcrafted structural features and spatial edges derived from three-dimensional coordinates were excluded, while RNA backbone connectivity and secondary-structure base-pair edges were retained. All configurations were trained on TR60+HARIBOSS using the same training settings and evaluated on the four independent benchmarks. The sequence-derived configuration achieved an MCC of 0.631 on RB9 and 0.345 on TE18, compared with 0.652 and 0.350, respectively, for the complete model incorporating handcrafted and tertiary-structure features. The performance difference was larger on TL12 and JL10. These results demonstrate that sequence-derived representations, together with RNA backbone and secondary-structure connectivity, can support binding-site prediction even when explicit tertiary-structure features are unavailable.

We next evaluated combinations of the RNA language-model representations to examine whether integrating complementary sequence-derived features could improve prediction performance (Table 4). Among the individual LMs, RNA-FM achieved the highest MCC across all four benchmarks. However, its predictions showed a precision–recall imbalance, with recall exceeding precision on each dataset, indicating a greater proportion of false-positive predictions. The other individual LMs, RNABERT and ERNIE-RNA, showed substantially lower MCC values, suggesting limited predictive performance when used independently in the present model architecture. To improve the overall prediction performance, we evaluated pairwise combinations of the LM representations. Combinations involving RNA-FM generally achieved higher MCC values than those involving only RNABERT and ERNIE-RNA. In particular, RNABERT+RNA-FM provided a favourable overall performance, achieving MCC values of 0.634, 0.345, 0.602, and 0.351 on RB9, TE18, TL12, and JL10, respectively. Notably, this combination also provided a more balanced precision–recall profile than RNA-FM alone on several benchmarks. Adding all three LM representations did not further improve performance consistently. These results identify RNABERT+RNA-FM as the most favourable LM combination for sequence-derived binding-site prediction in the absence of handcrafted and explicit tertiary-structure features.

**Table 4.** Performance of different RNA language-model representations and their combinations on independent benchmark datasets.

| LMs |  | ERNIE | RNA-FM | RNABERT | ENIE-RNAFM | RNABERT-ERNIE | **RNABERT-RNAFM** | ALL three LMs |
|---|---|---|---|---|---|---|---|---|
| **RB9 test set** | MCC | 0.3081 | 0.6220 | 0.0960 | 0.6311 | 0.2839 | **0.6341** | 0.6094 |
|  | Precision | 0.5074 | 0.6907 | 0.4516 | **0.76** | 0.6666 | 0.7333 | 0.7343 |
|  | Recall | 0.5112 | **0.7894** | 0.1052 | 0.7142 | 0.24060 | 0.7443 | 0.7067 |
| **TE18 test set** | MCC | 0.1137 | 0.3494 | 0.1128 | 0.2756 | 0.2822 | **0.3448** | 0.2679 |
|  | Precision | 0.4246 | 0.5578 | 0.4838 | 0.5588 | **0.6760** | 0.5730 | 0.5405 |
|  | Recall | 0.3483 | **0.5955** | 0.1685 | 0.4269 | 0.2696 | 0.5505 | 0.4494 |
| **TL12 test set** | MCC | 0.3116 | 0.5997 | 0.2338 | 0.5805 | 0.2584 | **0.6024** | 0.5714 |
|  | Precision | 0.6239 | 0.8091 | 0.6792 | 0.8468 | 0.7291 | **0.8584** | 0.8571 |
|  | Recall | 0.4649 | **0.6751** | 0.2292 | 0.5987 | 0.2229 | 0.6178 | 0.5732 |
| **JL10 test set** | MCC | 0.2260 | 0.3040 | 0.1635 | 0.3403 | 0.1991 | 0.3507 | **0.3537** |
|  | Precision | 0.5290 | 0.5173 | 0.5846 | 0.5855 | **0.6938** | 0.5911 | 0.6028 |
|  | Recall | 0.3408 | 0.5580 | 0.1423 | 0.4868 | 0.1273 | **0.4981** | 0.4831 |

### 3.4 Feature Ablation Study

A feature ablation study was performed to evaluate the contributions of geometric features and solvent-accessible surface area (SASA) to RSMBind performance. RNA-FM was used as the language-model representation in this analysis, as it provided the most favourable overall performance among the evaluated language models and their combinations with geometric features. The results for other language-model configurations are provided in Supplementary Table S5. Four feature configurations were compared by including or excluding geometric features and SASA while keeping the model architecture and training settings unchanged. The inclusion of geometric features consistently improved MCC across all four independent benchmark datasets. Without SASA, adding geometric features increased MCC from 0.6311 to 0.6523 on RB9, 0.3448 to 0.3500 on TE18, 0.6024 to 0.6555 on TL12, and 0.3507 to 0.3982 on JL10. Similar improvements were observed when SASA was included. These results indicate that geometric descriptors provide additional structural information beyond the sequence-derived representation from RNA-FM. In contrast, SASA did not provide a consistent improvement across the datasets. When combined with geometric features and RNA-FM, SASA reduced MCC on RB9, TE18, and TL12, while a marginal improvement was observed on JL10 (0.3982 to 0.4063). Without geometric features, adding SASA reduced performance across all four datasets. Based on these results, geometric features combined with RNA-FM, without SASA, were selected for the final RSMBind model. This configuration provided the strongest overall performance across the four benchmark datasets.

### 3.5. Model Analysis and Hyperparameter Exploration

A series of experiments was conducted during model development to determine the graph architecture, graph representation, and optimization settings for nucleotide-level RNA binding-site prediction. The investigated factors included graph convolution architecture, spatial edge attribute encodings, spatial distance cutoff, positive-class weighting, and loss function. We first compared different graph convolution architectures, including GraphConv, RGCNConv (Relational Graph Convolutional Network), and GATv2Conv (Graph Attention Network version 2). GATv2Conv was selected for the final model based on its ability to incorporate heterogeneous edge relationships and continuous spatial distance information. We further evaluated different graph representations by examining the contribution of edge-type information and alternative spatial edge attribute encodings. The spatial distance cutoff was also investigated to determine an appropriate neighborhood size for capturing spatial relationships between nucleotides. The optimization settings were evaluated by comparing different positive-class weights and loss functions. Positive-class weights of 1.5, 2.0, 2.5, and 3.0 were examined to account for class imbalance. Binary cross-entropy (BCE) and focal loss were also compared to assess their suitability for the training data. Based on these experiments, the final configuration employed GATv2Conv with edge-type information, normalized spatial distance attributes, an 8 Å spatial cutoff, a positive-class weight of 1.5, and BCE loss. Detailed experimental comparisons are provided in Supplementary Tables S6– S10.

### 3.6. Discussion

#### 3.6.1. Analysis of model performance across RNA structural regions

To examine the relationship between RNA structural context and small-molecule binding, nucleotides were broadly classified into stem and loop regions. Stem regions comprise paired nucleotides, whereas loop regions comprise unpaired nucleotides, including hairpins, bulges, internal loops, and junctions. This classification was used to provide an overall comparison between paired and unpaired regions without further subdividing the loop categories. In the training set, comprising 24,107 nucleotides from 358 RNA structures, 58.29% of the nucleotides were located in stem regions and 41.70% in loop regions. Despite their lower overall abundance, loop regions contained a higher proportion of binding nucleotides (32.97%) than stem regions (25.30%). This pattern was also observed in the independent test sets, with the proportion of binding nucleotides in loop regions exceeding that in stem regions across TE18, RB9, TL12, and JL10. These observations indicate an association between loop regions and small-molecule binding in the evaluated datasets, although binding nucleotides were also frequently present in stem regions (Figure 2). We further evaluated the prediction performance separately for stem and loop regions across the four independent test sets (Table 6). The model consistently achieved higher precision, recall, and F1-scores for loop regions than for stem regions. The difference was particularly evident in RB9, where the F1-score was 0.816 for loop regions compared with 0.691 for stem regions. A similar pattern was observed in TE18, although the performance in both regions was lower than that in RB9. These results suggest that the model identifies loop-associated binding sites more reliably, whereas prediction of binding nucleotides in stem regions remains comparatively challenging.

**Table 5.** Ablation study evaluating the contributions of geometric features and solvent-accessible surface area (SASA) using RNA LMs representations on four independent benchmark datasets. The LMs column indicates whether RNA-FM was included in the feature configuration.

| Geometry | LMs | SASA | RB9 test set |  |  | TE18 test set |  |  | TL12 test set |  |  | JL10 test set |  |  |
|---|---|---|---|---|---|---|---|---|---|---|---|---|---|---|
|  |  |  | MCC | Precision | Recall | MCC | Precision | Recall | MCC | Precision | Recall | MCC | Precision | Recall |
| Yes | NO | NO | 0.4557 | 0.5670 | 0.6992 | 0.2817 | 0.5074 | 0.5730 | 0.3208 | 0.5906 | 0.560 | 0.1964 | 0.4945 | 0.3408 |
| **YES** | **YES** | **NO** | **0.6523** | **0.7656** | **0.7368** | **0.3500** | **0.5808** | **0.5449** | **0.6555** | **0.8760** | **0.6751** | **0.3982** | **0.6614** | **0.4756** |
| YES | NO | YES | 0.4618 | 0.6230 | 0.6090 | 0.2525 | 0.5663 | 0.3595 | 0.2872 | 0.6106 | 0.439 | 0.2277 | 0.5454 | 0.3146 |
| YES | YES | YES | 0.5677 | 0.7154 | 0.6616 | 0.2536 | 0.5306 | 0.4382 | 0.6188 | 0.9029 | 0.5923 | 0.4063 | 0.6719 | 0.4756 |
| NO | YES | NO | 0.6311 | 0.7333 | 0.7443 | 0.3448 | 0.5730 | 0.5505 | 0.6024 | 0.8584 | 0.6178 | 0.3507 | 0.5911 | 0.4981 |

**Table 6.**
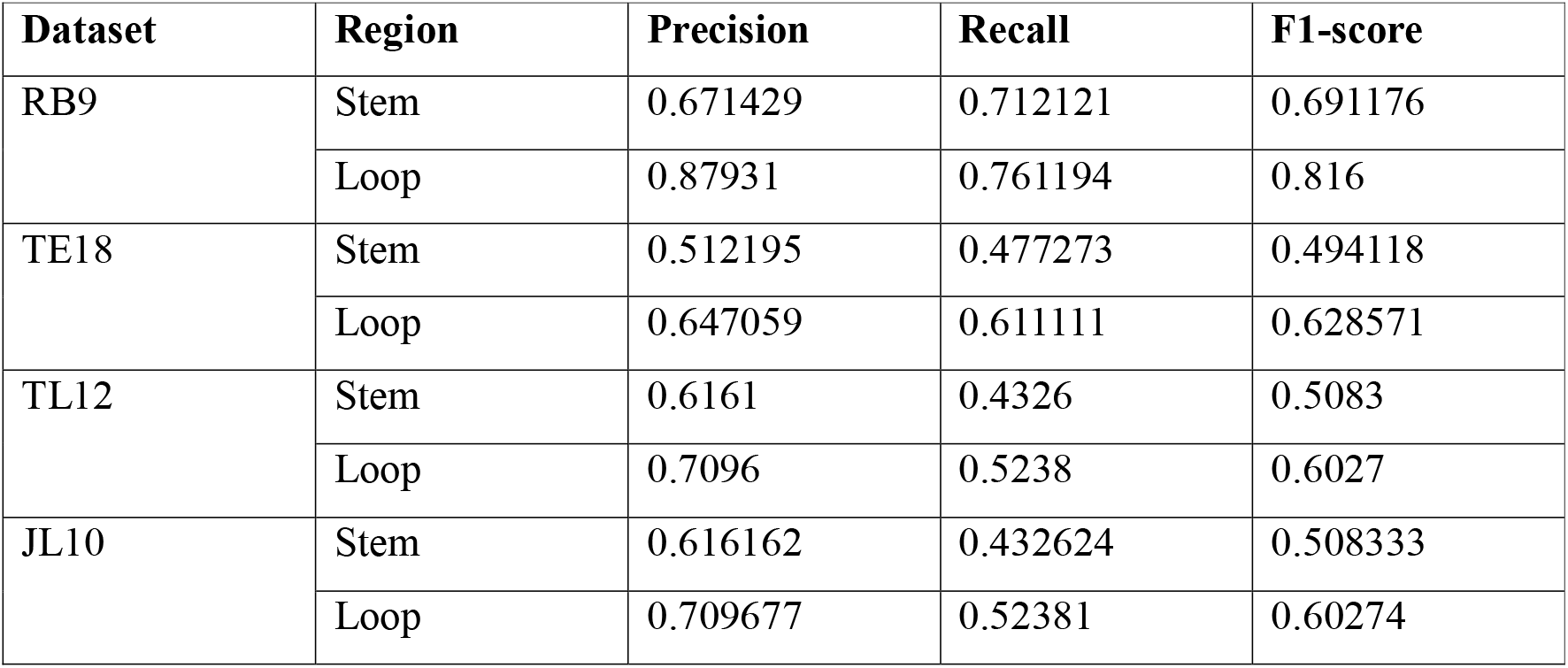
Prediction performance of RSMBind in stem and loop regions across the independent test sets.

**Figure 2:**
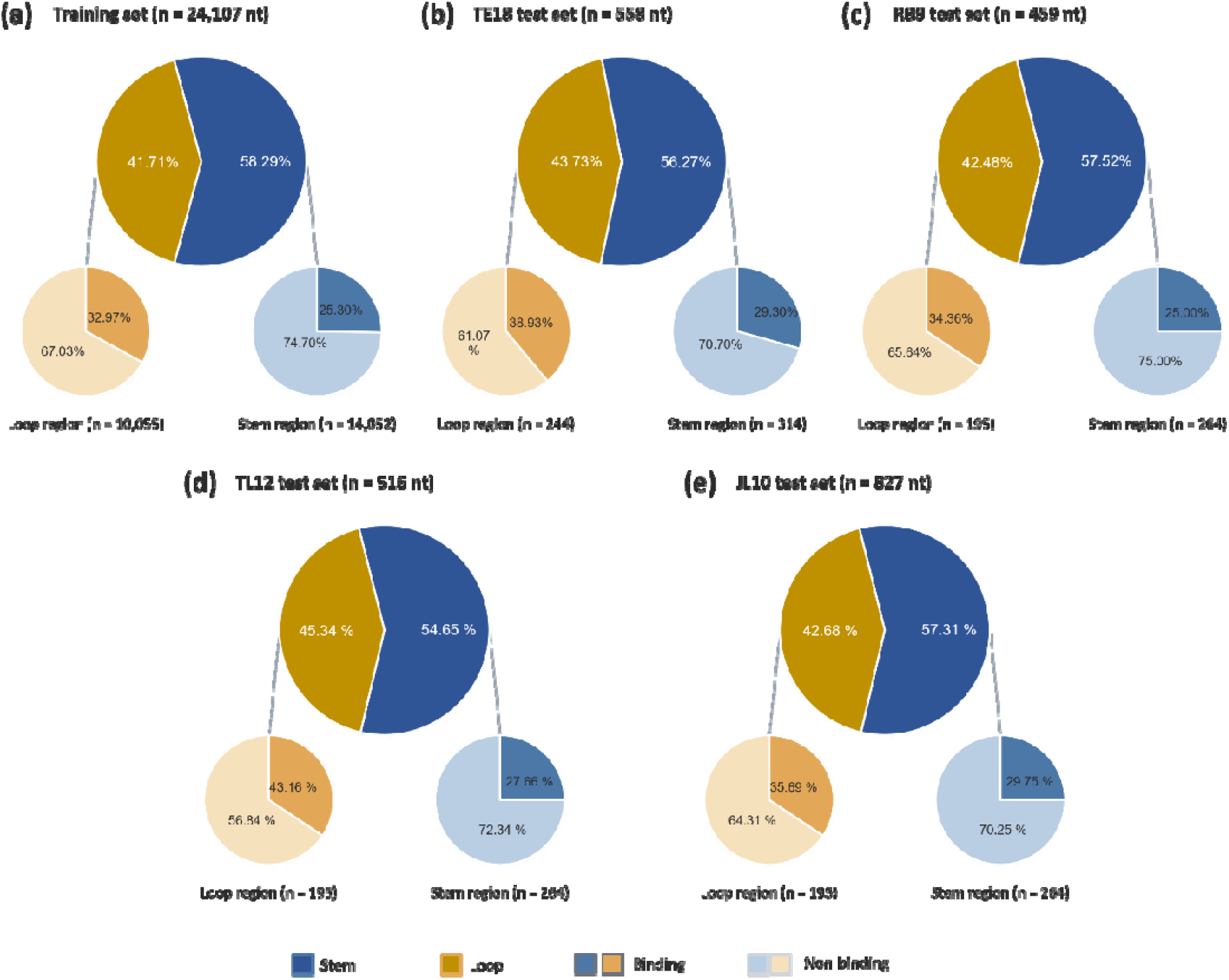
Region-wise distribution of binding and non-binding nucleotides in (a) the training set and the independent test sets: (b) TE18, (c) RB9, (d) TL12, and (e) JL10. The central pie chart shows the overall proportion of nucleotides in stem and loop regions, while the surrounding pie charts show the proportions of binding and non-binding nucleotides within each region.

To examine prediction differences at the individual structure level, we compared experimentally identified and model-predicted binding nucleotides in three representative complexes from the TE18 dataset (Figure 3). In 1F1T_A, containing 38 nucleotides and 11 binding nucleotides, the model correctly identified 10 binding positions, with one missed binding position and one false-positive prediction. In 1FMN_A (35 nucleotides and six binding positions), the model identified three binding nucleotides but also produced several false-positive predictions. In contrast, none of the six experimentally identified binding nucleotides in 4F8U_B (21 nucleotides) was correctly predicted. The secondary structures show that several binding nucleotides in 1F1T_A and 1FMN_A are located in loop regions or near stem–loop boundaries, whereas the binding region in 4F8U_B is predominantly stem-associated, with only a small bulge. This observation suggests that the model may have limitations in identifying binding sites within extended stem regions. The corresponding tertiary structures (Figure S2) provide additional spatial context for these RNA–ligand interactions. Together, these examples illustrate how prediction performance can vary with the structural context of binding nucleotides, complementing the region-wise performance analysis. To further investigate these prediction differences, we examined the three complexes using molecular dynamics (MD) simulations, focusing on ligand proximity and RNA structural dynamics.

**Figure 3.**
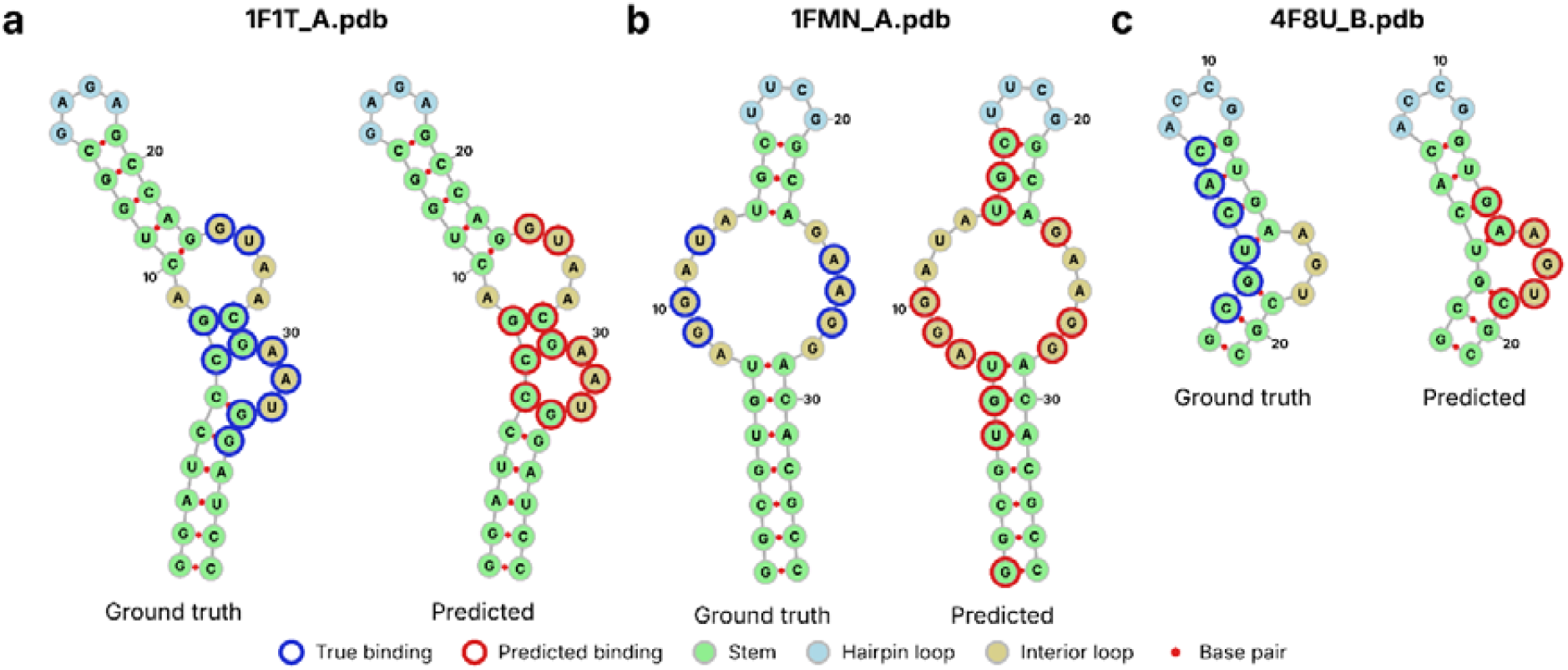
Comparison of experimentally identified and model-predicted binding nucleotides in the secondary structures of three representative RNA–small-molecule complexes from the TE18 dataset: (a) 1F1T_A, (b) 1FMN_A, and (c) 4F8U_B. Blue circles indicate experimentally identified binding nucleotides, while red circles indicate model-predicted binding nucleotides. Nucleotide colours represent the corresponding secondary-structure regions.

#### 3.6.2. Case-based analysis of model behaviour using MD simulations

To examine the relationship between model predictions and dynamic RNA–ligand proximity, three representative complexes from the TE18 dataset were selected: 1F1T_A, 1FMN_A and 4F8U_B, representing good, partial and poor prediction cases, respectively (Figure 4). The MD systems were prepared using CHARMM-GUI and simulated for 100 ns using GROMACS. For each nucleotide, the minimum distance between its atoms and the ligand heavy atoms was calculated for every trajectory frame. Three measures were considered: the trajectory-wide median distance, the closest approach and contact occupancy (percentage of frames with a distance below the 4 Å annotation cutoff). For 1F1T_A, Sr^2^□ was excluded from the MD system, and the analysis therefore represents RNA–ROS proximity only. 1F1T_A demonstrated the strongest agreement between model predictions and MD-derived ligand proximity. The model correctly identified all seven ROS-associated nucleotides (recall = 1.000; precision = 0.875). These nucleotides maintained close proximity to ROS, with median distances of 3.3–4.3 Å. Six of the seven exhibited contact occupancy above 86%, while G24 showed a comparatively lower occupancy of 41.8%. This agreement indicates that the model successfully identified the RNA region associated with persistent ligand proximity in the simulated complex. The additional predicted nucleotide, C6, was also located near the ligand (median distance = 6.4 Å), although its contact occupancy was below 0.1%. Thus, the predictions were largely consistent with the experimentally annotated binding region and its dynamic proximity to ROS.

**Figure 4.**
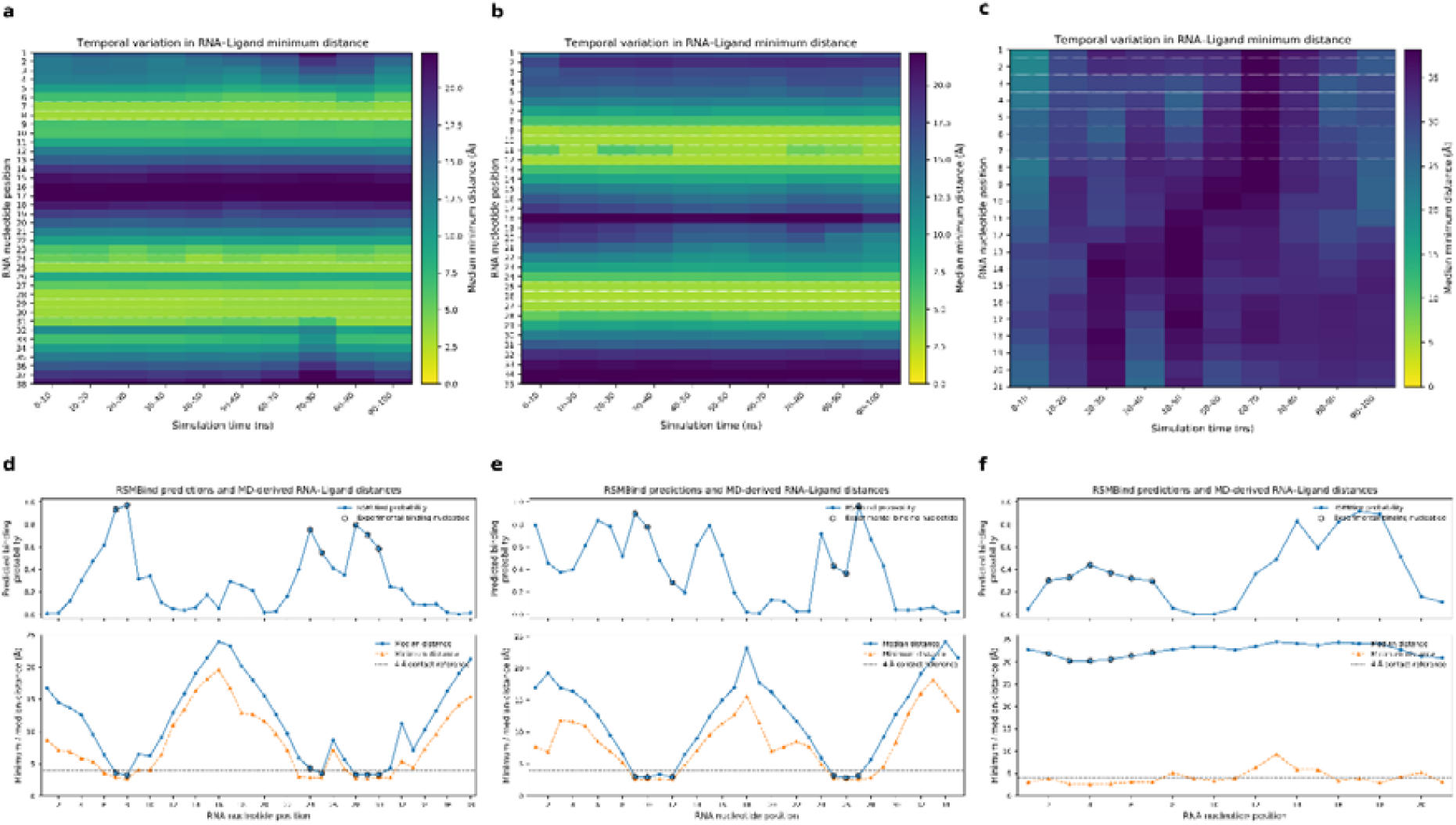
Case-based comparison of RSMBind predictions and MD-derived RNA–ligand proximity in three representative RNA–ligand complexes. (a–c) Heatmaps showing the temporal variation in nucleotide-wise minimum RNA–ligand distances over the 100 ns MD trajectories for (a) 1F1T_A, (b) 1FMN_A, and (c) 4F8U_B. (d–f) Corresponding nucleotide-wise RSMBind binding probabilities (upper panels) and trajectory-wide median and minimum RNA–ligand distances (lower panels) for (d) 1F1T_A, (e) 1FMN_A, and (f) 4F8U_B. Open circles indicate experimentally annotated binding nucleotides, and the dashed horizontal line in the lower panels represents the 4 Å distance threshold used for binding-site annotation.

1FMN_A illustrated a more challenging binding environment, with partial agreement between predictions and MD-derived proximity. The model identified three of the six experimentally annotated nucleotides (recall = 0.50; precision = 0.230). The detected nucleotides (G9, G10 and G27) showed high prediction probabilities (0.78–0.96) and persistent ligand contact. Interestingly, the three undetected experimental sites also maintained close proximity to the ligand, with median distances of 2.9–3.2 Å and contact occupancy of 99.7–100%, despite their relatively lower prediction probabilities (0.29–0.43). This indicates that the model did not fully capture the ligand-associated region, even when the corresponding nucleotides remained in close contact during the simulation. Nevertheless, several additional predicted nucleotides were located near the annotated binding region. In particular, A8, G24 and G28 were sequence-adjacent to experimentally annotated positions, with G28 exhibiting ligand contact in 6.5% of frames. These observations suggest that the model can recognize features associated with the broader binding environment, although distinguishing individual interacting nucleotides remains challenging in some cases. 4F8U_B provided an example of a more substantial limitation in model prediction. None of the six experimentally annotated binding nucleotides (C2–C7) were identified, and their prediction probabilities ranged from 0.30 to 0.44. Unlike the other two complexes, these nucleotides showed large median distances from the ligand (30.2–32.0 Å), with contact occupancy of only 0.1–1.6%. Although their closest approaches occasionally fell below 4 Å, such contacts were rare and transient. Furthermore, the high-probability predictions (G14–C19; 0.51–0.92) did not show comparable ligand proximity, with median distances of 32.7–34.3 Å. Thus, the model predictions did not correspond to the experimentally annotated interaction region or to the ligand proximity observed during the simulated trajectory. This case highlights a potential limitation of predicting binding sites from static structural features, particularly when the experimentally observed binding arrangement is not maintained in the simulated conformational ensemble. However, the lack of persistent contacts in this trajectory does not establish the absence of the experimentally observed binding interaction.

Overall, the three cases illustrate different levels of agreement between structure-based predictions and dynamic RNA–ligand proximity. The strong agreement in 1F1T_A demonstrates the ability of the model to identify nucleotides associated with persistent ligand proximity. The partial agreement in 1FMN_A indicates that the model can capture important parts of a binding environment but may not identify all interacting nucleotides. In contrast, 4F8U_B illustrates a more challenging case in which the predicted sites and experimentally annotated binding region were not supported by the observed ligand proximity during simulation. Since RSMBind relies primarily on features derived from experimentally determined RNA structures, its predictions may be influenced by the structural representation of the binding environment. Incorporating conformational variability and MD-derived dynamic features could therefore provide complementary information for improving RNA– small-molecule binding-site prediction.

## 4. Conclusion

In this study, we introduced RSMBind, a graph attention-based deep learning framework for nucleotide-level prediction of RNA–small-molecule binding sites by integrating local three-dimensional geometric features with pretrained RNA language model representations. The framework represents RNA as a spatial graph, allowing graph attention to incorporate information from structurally related neighboring nucleotides while combining complementary sequence and geometric information. We systematically evaluated RNA-BERT, ERNIE-RNA and RNA-FM embeddings to investigate their contributions to binding-site prediction. Evaluation on four independent benchmark datasets (RB9, TE18, TL12 and JL10) demonstrated the predictive capability of RSMBind across different RNA sequences and structural contexts. Compared with the best previously reported MCC values, RSMBind achieved relative improvements of 17.1%, 7.2% and 16.0% on RB9, TL12 and JL10, respectively, while showing competitive performance on TE18. Ablation experiments further demonstrated the complementary contributions of pretrained RNA language model representations and local geometric descriptors to nucleotide-level prediction. Structural analysis revealed differences in prediction performance across RNA structural contexts, with loop-associated and structurally exposed binding sites generally being captured more effectively than those embedded within stem regions. The MD-based case studies provided additional insights into the relationship between model predictions and dynamic RNA–ligand proximity, highlighting both the agreement and limitations of structure-based predictions. Despite these promising results, limitations remain due to the limited availability of high-resolution RNA–small-molecule structures and the ability of the current geometric representation to capture complex three-dimensional binding environments. Future work will explore additional geometric descriptors related to surface organization and molecular accessibility, larger RNA language models, and conformationally informed representations to improve prediction robustness and generalization. Overall, RSMBind demonstrates the value of integrating pretrained RNA sequence representations with graph-based three-dimensional structural information and provides a foundation for developing more geometry-aware computational approaches for RNA–small-molecule binding-site prediction.

## Acknowledgment

We do hereby acknowledge the use of the Supercomputer facility Paramshakti of the Indian Institute of Technology Kharagpur.

## Authors

Nandita Sharma received the B.Tech. degree from the University of Burdwan and the M.Tech. degree from NIT Patna, India. She is currently pursuing a Ph.D. degree in the Centre for Computational and Data Sciences at IIT Kharagpur under the supervision of Dr. Pralay Mitra. Her research focuses on RNA structure and functional analysis.

Pralay Mitra received the B.Sc. (with Hons.) degree in physics, and the B.Tech. degree in computer science and engineering from the University of Calcutta, Kolkata, India, in 1999 and 2002, respectively, and the M.E. degree in computer science and engineering from the Indian Institute of Engineering Science and Technology, Shibpur, Howrah, India, (formerly Bengal Engineering College and Science University, Shibpur), in 2004, and the Ph.D. degree from the Indian Institute of Science, Bangalore, Bengaluru, India, in 2010. Since 2005, he is actively working on Bioinformatics and Computational Biology. He has developed a number of algorithms for protein-protein docking, predicting protein assembly from the crystal structures, and protein design. He was a Research Associate with the Indian Institute of Science, Bangalore from 2010 to 2011 and Research Fellow with the University of Michigan, Ann Arbor, MI, USA from 2011 to 2013. Currently, he is an Associate Professor with the Department of Computer Science and Engineering, Indian Institute of Technology Kharagpur, India.

## Authors Contribution

N.S.: Conceptualization, Data curation, Formal analysis, Investigation, Methodology, Software, Validation, Visualization, Writing – original draft, Writing – review and editing

P.M.: Conceptualization, Formal analysis, Investigation, Methodology, Resources, Supervision, Writing – original draft, Writing – review and editing

