## Supplementary material for "RNA Small-Molecule Binding-Site Prediction by Integrating RNA Language Models and Structural Geometry": Table S1, Table S2, Table S3, Table S4, Table S5, Table S6, Figure S1, Figure S2

Pralay Mitra, Ph.D.

Department of Computer Science and Engineering,

Indian Institute of Technology Kharagpur

West Bengal - 721302, India

ORCID ID: 0000-0003-4119-3788

**Table of Contents**

| **Section** | **Page No** |
| --- | --- |
| S1. Details of Benchmark dataset | 3 |
| S2. Feature distribution analysis | 6 |
| S3. Model ablation | 7 |
| S4. Model Architecture and Hyperparameter Exploration | 8 |

S1. **Details of Benchmark dataset**

**Table S1.** Sequence, Secondary structure of test set RB9 with Dot-Bracket representation.

| PDB_FILE | SEQUENCE | Secondary Structure |
| --- | --- | --- |
| 1NEM_A.pdb | GGACUGGGCGAGAAGUUUAGUCC | (((((((((.....))))))))) |
| 1PBR_A.pdb | GGCGUCACACCUUCGGGUGAAGUCGCC | ((((.(.((((....))))..).)))) |
| 1Q8N_A.pdb | GGAUCCCGACUGGCGAGAGCCAGGUAACGAAUGGAUCC | ((((((((.(((((....)))))....))...)))))) |
| 1UUI_B.pdb | GGCAGAUCUGAGCCUGGGAGCUCUCUGCC | ((((((...((((......)))))))))) |
| 1Y26_X.pdb | CGCUUCAUAUAAUCCUAAUGAUAUGGUUUGGGAGUUUCUACCAAGAGCCUUAAACUCUUGAUUAUGAAGUG | (((((((((...((((((.........))))))........((((((.......))))))..))))))))) |
| 2GDI_X.pdb | NGACUCGGGGUGCCCUUCUGCGUGAAGGCUGAGAAAUACCCGUAUCACCUGAUCUGGAUAAUGCCAGCGUAGGGAAGUUC | .((..((((..(((.(((.....)))))).........))))..)).((((..((((......))))..))))....... |
| 2TOB_A.pdb | ACUUGGUUUAGGUAAUGAGU | (((((...)))))....... |
| 3D2X_A.pdb | GGGACCAGGGGUGCUUGUUCACAGGCUGAGAAAGUCCCUUUGAACCUGAACAGGGUAAUGCCUGCGCAGGGAGUGUC | (((((.......(((((....))))).......)))))......((((..((((......))))..))))....... |
| 3GX2_A.pdb | GGCUUAUCAAGAGAGGUGGAGGGACUGGCCCGACGAAACCCGGCAACCAGAAAUGGUGCCAAUUCCUGCAGCGGAAACGUUGAAAGAUGAGCCG | ((((((((......((((..(((.....)))..)...))).((((.(((....)))))))........(((((....)))))...)))))))). |

**Table S2.** Sequence, Secondary structure of test set TE18 with Dot-Bracket representation.

| PDB_FILE | SEQUENCE | Secondary Structure |
| --- | --- | --- |
| 1DDY_A.pdb | GGAACCGGUGCGCAUAACCACCUCAGUGCGAGCAA | .......((.(((((..........))))).)).. |
| 1F1T_A.pdb | GGAUCCCGACUGGCGAGAGCCAGGUAACGAAUGGAUCC | ((((((((.(((((....)))))....))...)))))) |
| 1FMN_A.pdb | GGCGUGUAGGAUAUGCUUCGGCAGAAGGACACGCC | (((((((......(((....))).....))))))) |
| 1NEM_A.pdb | GGACUGGGCGAGAAGUUUAGUCC | (((((((((.....))))))))) |
| 1Q8N_A.pdb | GGAUCCCGACUGGCGAGAGCCAGGUAACGAAUGGAUCC | ((((((((.(((((....)))))....))...)))))) |
| 2JUK_A.pdb | GGCCUUCCCACAAGGGAAGGCC | (((((((((....))))))))) |
| 2MIS_A.pdb | GAGCUGCAGCACGAAAGUGACGGCUC | ((((((...(((....))).)))))) |
| 2PWT_A.pdb | GCGUCACACCGGUGAAGUCGC | (((((((....))))...))) |
| 2TOB_A.pdb | ACUUGGUUUAGGUAAUGAGU | (((((...)))))....... |
| 364D_C.pdb | CCCCAUGCGAGAGUAGGGAACUGCCAGGCAU | ....((((..(.((((....)))))..)))) |
| 379d_B.pdb | GGCCGAAACUCGUAAGAGUCACCAC | ........(((....)))....... |
| 430D_A.pdb | GGGUGCUCAGUACGAGAGGAACCGCANCC | .(((.(((.......)))..)))...... |
| 4F8U_B.pdb | GCGUCACACCGGUGAAGUCGC | (((((((....))))...))) |
| 4PQV_A.pdb | GGGUCAGAUCGGCGAAAGUCGCCACUUCGCCGAGGAGUGCAAUCUGUGAGGCCCCAGGAGGACUGGGU | ..........(((((...))))).....(((..(.((......)).)..)))(((((.....))))). |
| 4YAZ_R.pdb | NGUACACGACAAUACUAAACCAUCCGCGAGGAUGGGGCGGAAAGCCUAAGGGUCUCCCUGAGACAGCCGGGCUGCCGAAAUAUC | ...................((((((....))))))..(((..(((((....(((((...)))))....))))).)))....... |
| 5BJO_E.pdb | GGCGCGAGGAAGGAGGUCUGAGGAGGUCACUGCGCC | ((((((..((......(((....)))))..)))))) |
| 5V3F_A.pdb | GUGCGAAGGGACGGUGCGGAGAGGAGAGCAC | ((((.......................)))) |
| 6EZ0_A.pdb | GGAGCGGGGGUGUAAACCUAUCGCUCC | ((.(((..((....)).))))) |

**Table S3.** Sequence, Secondary structure of test set JL10 with Dot-Bracket representation.

| PDB_FILE | SEQUENCE | Secondary Structure |
| --- | --- | --- |
| 6WTR_A.pdb | NGUUGCCGAAUCCUUUACUUGGUACGGAGGAACCGCUUUUUGGGGUUAAUCUGCAGUGAAGCUGCAGUAGGGAUACCUUCUGUCCCGCACCCGACAGCUAACUCCGGAGGCAAUAAAGGAAGGAG | .....((...(((((((.(((.(.(((((.....(((..(((((((....(((((((...)))))))..((((((.....)))))))).))))).)))...)))))..).)))))))))).)).. |
| 6WZS_B.pdb | GUUAUAUGACUGACGGAACGUGGAAUUAACCACAUGAAGUAUACGCCGACCGUCUGGGCG | ((((......)))).....((((......))))..........((((.........)))) |
| 7EAF_A.pdb | GGCUUAUCAAGAGAGGGCGAGGGACUGGCCCGACGACCCCCGGCAACCAGAAAUGGUGCCAAUUCCUGCAGCGGAAACGUUGAAAGAUGAGCCG | ((((((((....(.(((((.(((.....)))..)).))).)((((.(((....)))))))........(((((....)))))...)))))))). |
| 7EQJ_B.pdb | GGGUGAUUAGCUCAGCUGGGAGAGCACCUCCCUUACAAGGAGGGGGUCGGCGGUUCGAUCCCGUCAUCACCCACC | (((((((.(((...)))(((((.....))))).......((.(((((((......))))))).)))))))))... |
| 7KGA_A.pdb | GGGGGCCAGAUGUCAUGUCUCUCAAGCCUAGGAGACACUAGACACUCUGGACUAUCGGUUAGAGGAAACCCCCCCAAAAAUGUAUAGGC | ((((((((((((((.(((((((........)))))))...)))).)))))......((((......))))))))).............. |
| 7KVU_G.pdb | NGGAAGAUACAAGGUGAGCCCAAUAAUAUGGUUUGGGUUAGGAUAGGAAGUAGAGCCUUAAACUCUCUAAGCGGUAUCUUCCC | .(((((((((......(((((((.........))))))).........((.((((.......)))))).....))))))))). |
| 7TZR_X.pdb | NGACUCGGGGUGCCCUUCUGCGUGAAGGCUGAGAAAUACCCGUAUCACCUGAUCUGGAUAAUGCCAGCGUAGGGAAGUUC | .((..((((..(((.(((.....)))))).........))))..)).((((..((((......))))..))))....... |
| 7UQ6_B.pdb | NCGGAAGUAGUUCAGUGGUAGAACACCGGGUGGAACCCCAGGAAACUGGCCCGGGUCGCGCGGGUUCGAGUCCCGUCUUCCGCUCC | .((((((..((((.......)))).(((((.......((((....))))))))).....(((((.......))))))))))).... |
| 7UZ0_A.pdb | NCGGAAGUAGUUCAGUGGUAGAACACCUACUGGCCCGAAAGGGUGGAACCCCAGUAGGGGGUCGCGGGUUCGAGUCCCGUCUUCCGC | .((((((.....((((((........))))))((((....)))).((.((((....)))).))(((((.......))))))))))). |
| 9BUN_C.pdb | GGACUCGGAAACGUGAAGGAGAGGCGCAAGGUUAACCGCCUCAGUCCA | (((((((....)).......((((((..........))))))))))). |

**Table S4.** Sequence, Secondary structure of test set TL12 with Dot-Bracket representation.

| PDB_FILE | SEQUENCE | Secondary Structure |
| --- | --- | --- |
| 7D12_A.pdb | GCAGCAGCUUCGGCAGCAGC | ((.((.((....)).)).)) |
| 7ELQ_A.pdb | GAGUAGAAGCGUUCAGCGGCCGAAAGGCCGCCCGGAAAUUGCUCC | ((((((...((....((((((....)))))).))....)))))). |
| 7EOG_A.pdb | GCGCACUGGCGCUGCGCCUUCGGGCGCCAAUCGUAGCGUGUCGGCGCC | ((((....((((((((((....))))).......)))))....)))). |
| 7L0Z_G.pdb | GGGAGUACGGUGAGGGUCGGGUCCAGUAGGUACGCCUACUGUUGAGUAGAGUGUGGGCUCCGUACUCCC | (((((((((((.(.(.((.(.(((((((((....)))))))..)).).)).).).)...)))))))))) |
| 7OA3_A.pdb | GCUAGCUGGAGGGGCGCCAGUUCGCUGGUGGUUGGGUGCGGUCGGCUAGCC | (((((((((....(((((...((......))...)))))..))))))))). |
| 7REX_A.pdb | UGUGGUUCGCAACCAUCCCACAUAAAAAAACUAG | (((((............)))))............ |
| 7RWR_A.pdb | GGAUACACAAGAGUGAUUGAAACUAAGUCUGUGUAUCC | (((((((((.((.(...........).))))))))))) |
| 7TD7_A.pdb | GCGACUCGGGGUGCCCCGUGAAGGCUGAGAAAUACCCGUAUCACCUGAUCUGGAUAAUGCCAGCGUAGGGAAGUCGCA | ((((((((((..(((.......))).........))))..((.((((..((((......))))..)))))))))))). |
| 8F0N_A.pdb | GCGCCGGUUAGGCAGUGGUGGGUGGUGUGGAGGAGUAGCUGUCCGGCGC | (((((((....(((((.....................)))))))))))) |
| 8FZA_A.pdb | UUGGGUUCCCUCACCCCAAUCAUAAAAAGG | ..((((......)))).............. |
| 8HB1_A.pdb | AGAGCGUUGCGUCCGAAAGUNGCC | .....((.((........)).)). |
| 8HB1_B.pdb | GCGACACGGCUCUUUAAAAACAAAAGGAGA | .........((((((........)))))). |

S2. **Feature distribution analysis**

To understand the structural characteristics associated with RNA–small-molecule binding sites, we performed a statistical analysis of nine handcrafted features on the training set. These features characterize three complementary aspects of RNA structure: surface properties (SASA), geometric properties, and local graph characteristics. The distributions of binding and non-binding nucleotides are shown in Figure S1. All feature values were standardized prior to visualization.

As shown in Figure S1, binding nucleotides generally exhibit lower SASA and centroid distance than non-binding nucleotides. In contrast, Pos Norm shows highly overlapping distributions between the two classes. The neighborhood-related features show differences in their distributions, with binding nucleotides generally exhibiting higher or broader values. Among the graph-based features, degree shows substantial overlap, whereas clustering coefficient shows a broader distribution toward higher values for binding nucleotides. Overall, these features capture complementary structural properties of RNA and provide information on the surface, spatial, and local structural environment of nucleotides.

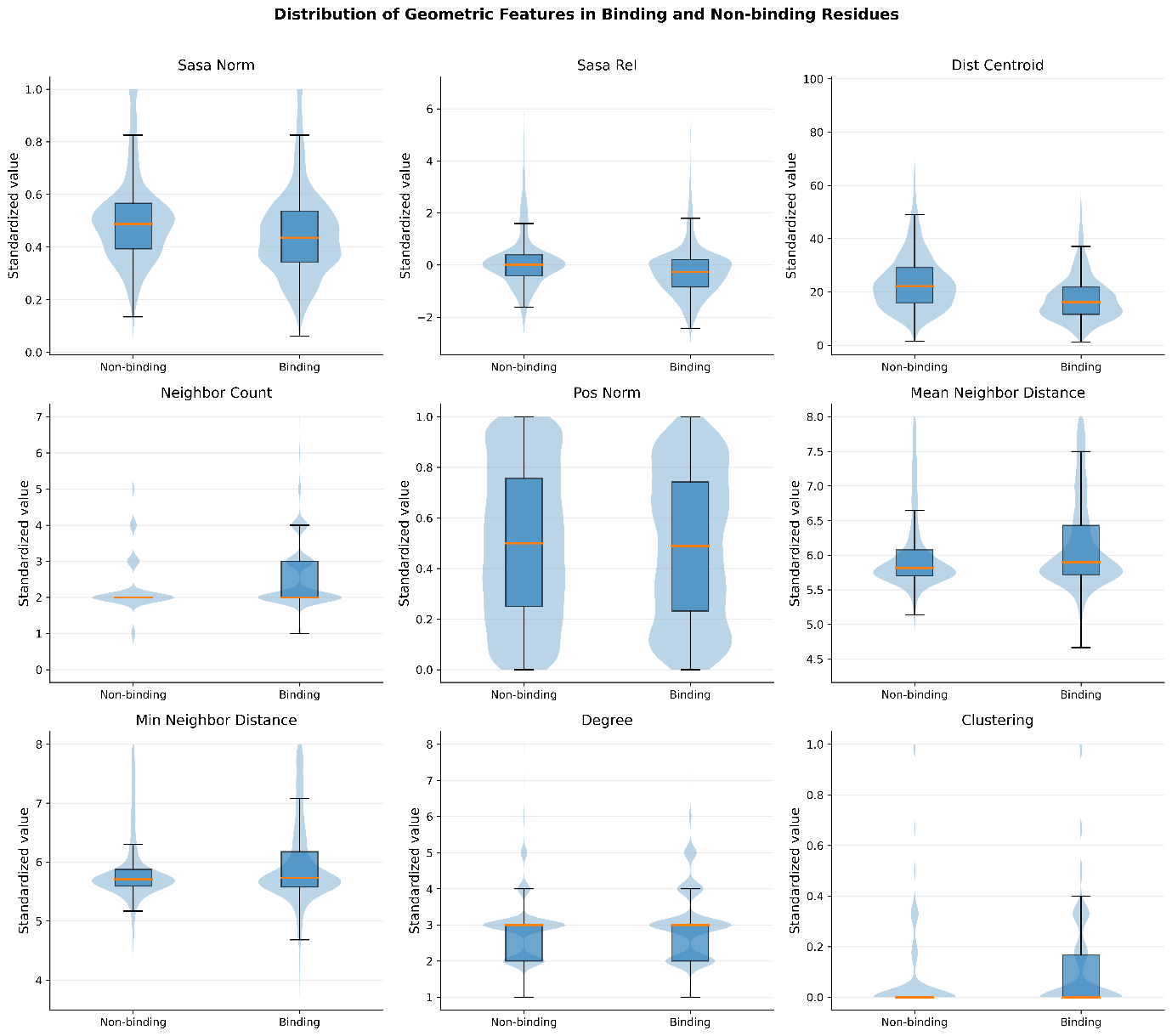

**Figure S1.** Violin and box plots showing the distributions of nine standardized handcrafted features for binding and non-binding nucleotides in the training set.

S3. **Model ablation**

To further examine the contributions of RNA language-model representations and solvent-accessible surface area (SASA), we evaluated different combinations of pretrained RNA language models, with and without SASA. In all configurations, the seven geometric and graph-topological descriptors described in the Methods were retained as the default structural features. The evaluated configurations included individual language models, pairwise combinations, all three language models, and a configuration without language-model features. This analysis allowed us to assess the contributions of different sequence-derived representations and SASA alongside the structural features. The results are summarized in Table S5.

**Table S5.** Performance of different RNA language-model configurations, with and without SASA, in combination with seven geometric and graph-topological descriptors on four independent benchmark datasets. The best result for each metric on each dataset is highlighted in bold.

| **LMs** | **SASA** | **RB9 test set** | | | **TE18 test set** | | | **TL12 test set** | | | **JL10 test set** | | |
| --- | --- | --- | --- | --- | --- | --- | --- | --- | --- | --- | --- | --- | --- |
|  |  | **MCC** | **Precision** | **Recall** | **MCC** | **Precision** | **Recall** | **MCC** | **Precision** | **Recall** | **MCC** | **Precision** | **Recall** |
| ALL | YES | 0.568 | 0.715 | 0.662 | 0.254 | 0.531 | 0.438 | 0.619 | 0.903 | 0.592 | 0.406 | 0.672 | 0.476 |
| ALL | NO | 0.640 | 0.707 | 0.797 | 0.353 | 0.543 | 0.640 | 0.606 | 0.836 | 0.650 | 0.387 | 0.640 | 0.487 |
| ERNIE | YES | 0.458 | 0.597 | 0.647 | 0.248 | 0.532 | 0.421 | 0.428 | 0.684 | 0.580 | 0.355 | 0.618 | 0.461 |
| ERNIE | NO | 0.449 | 0.617 | 0.594 | 0.307 | 0.585 | 0.444 | 0.401 | 0.692 | 0.516 | 0.331 | 0.618 | 0.412 |
| RNAFM | YES | 0.541 | 0.662 | 0.692 | 0.287 | 0.548 | 0.478 | 0.641 | 0.879 | 0.650 | 0.351 | 0.595 | 0.491 |
| **RNAFM** | **NO** | **0.652** | **0.766** | **0.737** | **0.350** | **0.581** | **0.545** | **0.656** | **0.876** | **0.675** | **0.398** | **0.661** | **0.476** |
| RNABERT | YES | 0.249 | 0.636 | 0.211 | 0.235 | 0.661 | 0.208 | 0.339 | 0.774 | 0.306 | 0.251 | 0.746 | 0.165 |
| RNABERT | NO | 0.285 | 0.660 | 0.248 | 0.257 | 0.657 | 0.247 | 0.303 | 0.730 | 0.293 | 0.266 | 0.776 | 0.169 |
| ERNIE-RNAFM | YES | 0.571 | 0.709 | 0.677 | 0.300 | 0.569 | 0.461 | 0.636 | 0.913 | 0.605 | 0.370 | 0.648 | 0.442 |
| ERNIE-RNAFM | NO | 0.602 | 0.678 | 0.774 | 0.344 | 0.541 | 0.624 | 0.622 | 0.810 | 0.707 | 0.343 | 0.569 | 0.524 |
| RNABERT-RNAFM | YES | 0.599 | 0.700 | 0.737 | 0.339 | 0.557 | 0.573 | 0.617 | 0.844 | 0.656 | 0.394 | 0.652 | 0.483 |
| RNABERT-RNAFM | NO | 0.560 | 0.689 | 0.684 | 0.293 | 0.556 | 0.472 | 0.609 | 0.873 | 0.611 | 0.376 | 0.636 | 0.472 |
| ERNIE-RNABERT | YES | 0.426 | 0.690 | 0.451 | 0.255 | 0.594 | 0.320 | 0.416 | 0.745 | 0.465 | 0.322 | 0.683 | 0.315 |
| ERNIE-RNABERT | NO | 0.331 | 0.576 | 0.429 | 0.278 | 0.598 | 0.360 | 0.339 | 0.684 | 0.414 | 0.286 | 0.640 | 0.300 |
| NO LMs | YES | 0.462 | 0.623 | 0.609 | 0.253 | 0.566 | 0.360 | 0.287 | 0.611 | 0.439 | 0.228 | 0.545 | 0.315 |
| NO LMs | NO | 0.456 | 0.567 | 0.699 | 0.282 | 0.507 | 0.573 | 0.321 | 0.591 | 0.561 | 0.196 | 0.495 | 0.341 |

S4. **Model Architecture and Hyperparameter Exploration**

Several experiments were conducted to examine the effects of graph architecture, graph representation, and optimization settings on model performance. The comparisons were performed using the training and validation datasets, while the independent benchmark datasets were used for model evaluation. GraphConv, RGCNConv, and GATv2Conv were compared to evaluate different neighborhood aggregation mechanisms. The comparison included five-fold validation performance. GATv2Conv was retained for the final model based on its suitability for incorporating both categorical edge types and continuous spatial distance attributes (Table S6).

**Table S6.** Comparison of graph convolution architectures

| Graph architecture | Validation MCC | Validation AUROC | Validation F1 |
| --- | --- | --- | --- |
| GraphConv | 0.5469 ± 0.0167 | 0.8295 ± 0.0107 | 0.6568 ± 0.0102 |
| RGCNConv | 0.5470 ± 0.0157 | 0.8307 ± 0.0072 | 0.6495 ± 0.0108 |
| GATv2Conv | 0.5696 ± 0.0109 | 0.8447 ± 0.0109 | 0.6599 ± 0.0087 |

The effects of edge-type information and spatial edge attributes were investigated. Different spatial distance encodings were compared, including exponential distance transformations and normalized distances (Table S7). Spatial cutoffs of 6, 8, and 10 Å were also evaluated. The selected graph representation retained edge-type information and normalized spatial distances, with an 8 Å spatial cutoff (Table S8). Positive-class weights of 1.5, 2.0, 2.5, and 3.0 were evaluated to examine their effects on classification performance (Table S9). BCE and focal loss $\left( \left( \gamma=2 \right) \right)$were also compared. Based on the validation results, a positive-class weight of 1.5 and BCE loss were selected for the final model (Table S10).

**Table S7.** Evaluation of Spatial Edge Attribute Encodings

| Spatial edge attribute encoding | MCC | AUROC | F1-score |
| --- | --- | --- | --- |
| $\exp\left( -d/8 \right)$ | 0.5376 | 0.8307 | 0.6412 |
| $d_{\mathrm{norm}}=d/8$ | 0.5696 | 0.8447 | 0.6599 |
| $\exp\left( -d/2 \right)$ | 0.5230 | 0.8199 | 0.6219 |

**Table S8.** Evaluation of Spatial Distance Cutoffs

| Spatial cutoff | MCC | AUROC | F1-score |
| --- | --- | --- | --- |
| 6 Å | 0.5370 | 0.8290 | 0.6511 |
| 8 Å | 0.5696 | 0.8447 | 0.6599 |
| 10 Å | 0.5215 | 0.8201 | 0.6199 |

**Table S9.** Effect of Positive-Class Weighting on Model Performance

| Positive-class weight | MCC | AUROC | F1-score |
| --- | --- | --- | --- |
| 1.5 | 0.5696 | 0.8447 | 0.6599 |
| 2.0 | 0.5540 | 0.8364 | 0.6578 |
| 2.5 | 0.5359 | 0.8306 | 0.6477 |
| 3.0 | 0.5402 | 0.8345 | 0.6561 |

**Table S10.** Comparison of Loss Functions

| Loss function | MCC | AUROC | F1-score |
| --- | --- | --- | --- |
| BCE | 0.5696 | 0.8447 | 0.6599 |
| Focal loss | 0.5272 | 0.8235 | 0.6235 |

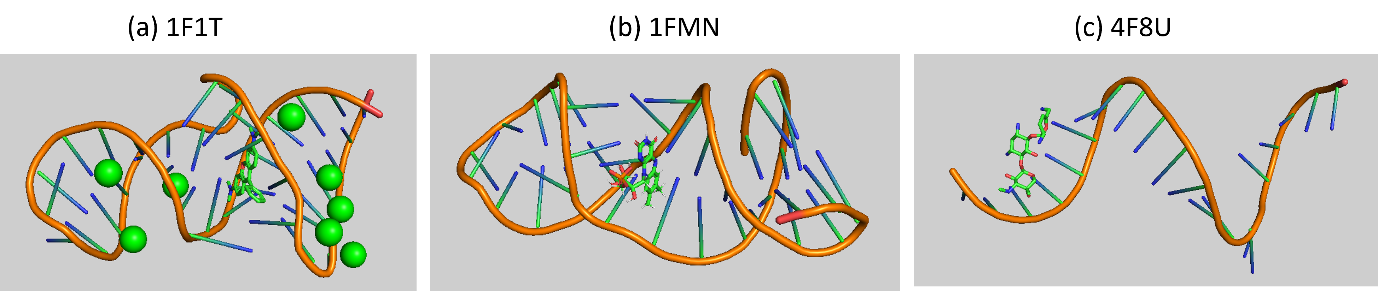

**Figure S2.** Tertiary-structure representations of the three representative RNA–small-molecule complexes from the TE18 dataset: (a) 1F1T_A, (b) 1FMN_A, and (c) 4F8U_B. RNA backbones are shown in orange, and the bound small molecules are represented as sticks. Green spheres in 1F1T_A represent the associated metal ions.
